# Whole-genome duplications and the long-term evolution of gene regulatory networks in angiosperms

**DOI:** 10.1101/2023.03.13.532351

**Authors:** Fabricio Almeida-Silva, Yves Van de Peer

**Affiliations:** Department of Plant Biotechnology and Bioinformatics, Ghent University, 9052 Ghent, Belgium; VIB Center for Plant Systems Biology, VIB, 9052 Ghent, Belgium; Centre for Microbial Ecology and Genomics, Department of Biochemistry, Genetics and Microbiology, University of Pretoria, Pretoria 0028, South Africa; College of Horticulture, Academy for Advanced Interdisciplinary Studies, Nanjing Agricultural University, Nanjing, China

**Keywords:** systems biology, genome evolution, polyploidy, network evolution, bioinformatics

## Abstract

Angiosperms have a complex history of whole-genome duplications (WGDs), with varying numbers and ages of WGD events across clades. These WGDs have greatly affected the composition of plant genomes due to the biased retention of genes belonging to certain functional categories following their duplication. In particular, regulatory genes and genes encoding proteins that act in multiprotein complexes have been retained in excess following WGD. Here, we inferred protein-protein interaction (PPI) networks and gene regulatory networks (GRNs) for seven well-characterized angiosperm species and explored the impact of both WGD and small-scale duplications (SSD) in network topology by analyzing changes in frequency of network motifs. We found that PPI networks are enriched in WGD-derived genes associated with dosage-sensitive intricate systems, and strong selection pressures constrain the divergence of WGD-derived genes at the sequence and PPI levels. WGD-derived genes in network motifs are mostly associated with dosage-sensitive processes, such as regulation of transcription and cell cycle, translation, photosynthesis, and carbon metabolism, while SSD-derived genes in motifs are associated with response to biotic and abiotic stress. Recent polyploids have higher motif frequencies than ancient polyploids, while WGD-derived network motifs tend to be disrupted on the longer term. Our findings demonstrate that both WGD and SSD have contributed to the evolution of angiosperm GRNs, but in different ways, with WGD events likely having a more significant impact on the short-term evolution of polyploids.

## Introduction

Whole-genome duplications (WGD) are a key source of extra genetic material for evolution to work with (Ohno 1970), and they have occurred independently in multiple branches of the tree of life. For instance, vertebrates have experienced two rounds of WGD early in their evolution (Dehal and Boore 2005; Van de Peer et al. 2010; Nakatani et al. 2021), with an additional WGD event shared by all teleosts (Amores et al. 1998; Taylor et al. 2003). The genomes of several yeasts have also undergone ancient WGD events (Wolfe and Shields 1997). However, WGD events have been most abundant in plants, and they are thought to have contributed to the radiation of important plant families (Schranz et al. 2012; Tank et al. 2015; Landis et al. 2018), to increased diversity (Ren et al. 2018), and to survival in stressful times (Van de Peer et al. 2017; Van de Peer et al. 2021).

After a WGD event, survival and establishment of newly formed polyploids is challenging due to the detrimental effects of doubling the entire set of chromosomes, which leads to e.g., genomic shock and reduced fertility (Comai 2005; Woodhouse et al. 2010). Polyploids that survive a WGD event typically undergo a rediploidization process that leads to genome fractionation (i.e., loss of functional DNA sequences), probably because of functional redundancy (De Smet et al. 2013; Li et al. 2016). During the process of rediploidization and fractionation, it has been observed that genes encoding proteins involved in intricately connected systems, such as transcription factors, kinases, and members of multiprotein complexes, tend to be retained more often than other genes after WGD (Blanc and Wolfe 2004; Maere et al. 2005; Freeling 2009). To date, this biased retention of particular gene classes is best explained by the gene balance hypothesis, which states that some gene families are dosage sensitive, hence their retention preserves stoichiometric balance (Freeling and Thomas 2006; Birchler and Veitia 2010). For the same reason, retention of such dosage-sensitive genes is usually selected against when these are duplicated in small-scale duplication (SSD) events, as stoichiometric balance is disrupted (Maere et al. 2005; Freeling 2009).

The preferential retention of transcription factors (TFs) following WGDs motivated researchers to explore the impact of WGD events on the evolution of the TF repertoires and gene regulatory networks (GRNs) (De Smet and Van de Peer 2012; Panchy et al. 2019; Moharana and Venancio 2020; Mottes et al. 2021; Gera et al. 2022). In particular, WGD can reorganize the topology of GRNs by creating novel network motifs, which are enriched subgraphs typically found in complex networks (Milo et al. 2002). Network motifs are simple genetic circuits regarded as the building blocks of biological networks, and the connections that occur within and between motifs result in robust regulatory interactions observed in complex biological systems (Milo et al. 2002; Burda et al. 2011). As motifs perform elementary regulatory functions, they have been positively selected during the evolution of biological networks (Ward and Thornton 2007; Mottes et al. 2021), and examples include V motifs (paralogous TFs regulating a shared target gene), lambda motifs (paralogous target genes regulated by a shared TF), and bifans (a pair of paralogous TFs that regulate the same pair of targets) (Mottes et al. 2021).

Analyzing motif frequencies can reveal the impact of gene and genome duplications in biological networks and thus evolutionary innovation or (increase in) biological complexity. For instance, in human GRNs, WGD events have increased regulatory redundancy (and, hence, increased mutational robustness), and generated complex combinations of motifs, which enhanced the performance of high-level functions such as signal integration and noise control (Mottes et al. 2021). Likewise, WGD events led to the enrichment of bifans in yeast, which have been shown to be important for signal processing and formation of modular network structures (Ward and Thornton 2007). However, little is known about the impact of WGD on the topology of gene regulatory networks in angiosperms. As angiosperms have a more diverse WGD background, both with respect to the number and age(s) of WGD events across clades, studying the evolution of GRN topologies will further our understanding on the significance of WGDs - versus SSDs - for the evolution of flowering plants.

Here, we explored the impact of WGD and SSD events on protein-protein interaction (PPI) networks and GRNs of seven angiosperm species with well-defined WGDs. We observe that PPI networks are enriched in WGD-derived genes associated with intricately connected molecular processes. We show that interacting WGD-derived gene pairs have lower rates of sequence divergence and higher preservation of interactions as compared to SSD-derived gene pairs. Besides, genes originating from recent WGD events have higher numbers of GRN motifs than genes from ancient WGD events, and recent polyploids generally have higher motif frequencies. WGD events are the main source of novel motifs for recent polyploids, suggesting an immediate selective advantage for polyploids under certain conditions. However, WGD-derived motifs are lost over time and after adaptation. Finally, WGD-derived genes in GRN motifs seem associated with dosage-sensitive processes, such as transcriptional regulation, cell cycle, histone modifications, flower development, and photosynthesis, while SSD-derived genes in motifs seem associated with response to biotic and abiotic stress.

## Materials and Methods

### Genomic data acquisition

We obtained genomic data for seven angiosperm species: *Glycine max, Arabidopsis thaliana, Solanum lycopersicum, Populus trichocarpa, Vitis vinifera, Zea mays*, and *Oryza sativa* (Table 1). Protein sequences, DNA sequences for coding regions, and genome annotations were downloaded from Ensembl Plants release 53 (Yates et al. 2022) (Supplementary Table S1). RNA-seq data from baseline experiments were downloaded from EBI’s Expression Atlas (Papatheodorou et al. 2020) (Supplementary Table S1). Whole-genome functional annotations (Gene Ontology terms and InterPro domains) were obtained from Ensembl’s BioMart using the R package biomaRt (Durinck et al. 2009).

**Table 1.** Included species and summary statistics. Genes, number of genes in each genome. Ks peaks, number of peaks identified in whole-paranome Ks distributions. WGD and SSD, number of WGD- and SSD-derived gene pairs. GRN and PPI, number of edges (in millions) in each gene regulatory network and protein-protein interaction network, respectively.

| Species | Genes | Ks peaks | WGD | SSD | GRN (M) | PPI (M) |
| --- | --- | --- | --- | --- | --- | --- |
| *Glycine max* | 55897 | 2 | 34570 | 83219 | 2.27 | 1.38 |
| *Arabidopsis thaliana* | 27628 | 2 | 4329 | 43154 | 2.50 | 0.55 |
| *Solanum lycopersicum* | 34429 | 2 | 3906 | 57664 | 1.34 | 0.52 |
| *Populus trichocarpa* | 41335 | 3 | 15400 | 63613 | 1.28 | 0.64 |
| *Vitis vinifera* | 35134 | 1 | 2860 | 62571 | 1.21 | 0.21 |
| *Zea mays* | 39756 | 3 | 8095 | 69611 | 2.05 | 0.71 |
| *Oryza sativa* | 35775 | 2 | 3435 | 49498 | 1.80 | 0.25 |

### Prediction and classification of transcription factors

The R package *planttfhunter* (https://github.com/almeidasilvaf/planttfhunter/) was used to predict TFs from protein sequences and classify them into families using the same classification scheme implemented in PlantTFDB (Tian et al. 2020). Briefly, profile hidden Markov models for DNA-binding domains and auxiliary domains were searched against the entire proteome using HMMER 3.0 (Finn et al. 2011), and TF families were identified based on the presence and number of signature domains.

### Identification of duplicated genes and calculation of substitution rates

Duplicated gene pairs were identified and classified into whole-genome duplication (WGD)-derived and small-scale duplication (SSD)-derived gene pairs using the R/Bioconductor package *doubletrouble* (https://bioconductor.org/packages/doubletrouble). In short, all-against-all intragenomic DIAMOND protein similarity searches (sensitive mode, e-value = 1e-10, top hits = 5) were performed to identify the entire set of paralogous gene pairs for each species. Paralogous gene pairs in the same syntenic blocks were classified as WGD-derived pairs, and all other pairs were classified as derived from SSD. Rates of synonymous and nonsynonymous substitutions per substitution site (Ks and Ka, respectively) were calculated using the pipeline described in (Qiao et al. 2019), which consists in performing pairwise alignments with MAFFT (Katoh and Standley 2013) followed by Ka and Ks calculation with KaKs_Calculator 2.0 using the MYN model (Wang et al. 2010). Peaks in whole-paranome Ks distributions were identified with the R package *doubletrouble* by fitting Gaussian mixture models. As Gaussian mixture models can overestimate the number of peaks in Ks distributions, we explicitly defined the number of peaks for each species based on literature evidence (Qiao et al. 2019).

### Protein-protein interaction and gene regulatory network inference

Protein-protein interaction (PPI) networks were obtained from the STRING database using a confidence threshold of 0.5 (Szklarczyk et al. 2021). Gene regulatory networks were inferred using the GENIE3 algorithm (Huynh-Thu et al. 2010) as implemented in the R package BioNERO (Almeida-Silva and Venancio 2022). GRNs inferred with GENIE3 were represented as a fully connected bipartite graph. To remove spurious edges, we used the function *grn_filter()* from BioNERO to rank edges and split each GRN in 20 subnetworks of increasing number of edges, where the 1^st^ subnetwork contains the 0.05 quantile and the 20^th^ subnetwork is the original fully connected graph. For each subnetwork, the function *grn_filter()* fitted the degree distribution to a scale-free topology. The largest subnetwork with an R^2^ > 0.75 (coefficient of determination for the scale-free topology fit) was selected as the optimal GRN (Supplementary Figure S1).

### Network topology analyses and functional enrichment

To ensure that the topology of the inferred GRNs and PPI networks resembled the topology of real-world biological networks, all networks were checked for scale-free topology fit using the function *check_SFT()* from the R package BioNERO (Almeida-Silva and Venancio 2022). Degree distributions for WGD- and SSD-derived gene pairs were calculated with the R package igraph (Csardi and Nepusz 2006) and visualized with the R package ggstatsplot (Patil 2021). Enrichment analyses were performed with the function *enrichment_analysis()* from the R package BioNERO. *P*-values from Fisher’s exact tests were adjusted for multiple comparisons with Benjamini-Hochberg correction.

### Identification of network motifs, significance assessment, and interaction similarity

To identify and analyze graph motifs, we developed an R package named *magrene*, which is available on Bioconductor (https://bioconductor.org/packages/magrene), and the source code of which is available on a GitHub repository (https://github.com/almeidasilvaf/magrene). This package can be used to identify five types of motifs containing paralogous gene pairs, namely V, PPI V, lambda, delta, and bifan motifs (Figure 1). For each species, we calculated the frequency of motifs containing WGD- and SSD-derived gene pairs. To account for the effect of ‘gene age’ on motif frequencies, we classified duplicated gene pairs into age groups based on Ks peaks using the function *split_pairs_by_peak()* from the *doubletrouble* package. Briefly, age groups were defined based on the mean ± 2 standard deviations relative to each Ks peak, and duplicate pairs (either WGD- or SSD-derived) with Ks values within a particular age group were assigned to it. Then, WGD vs SSD comparisons were only performed for paralogs belonging to the same age group. To assess the significance of the observed motif frequencies, we used the function *generate_nulls()* from *magrene* to generate 1,000 degree-preserving simulated networks through node label permutation and calculate motif frequencies in each permutation. Then, the function *calculate_Z()* was used to obtain Z-scores for the observed motif frequencies. The function *sd_similarity()* was used to calculate the interaction similarity between gene pairs, which is represented by the Sorensen-Dice similarity index:

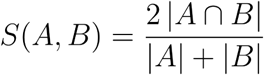

where A and B are the interacting partners of nodes A and B, respectively.

**Fig. 1.**
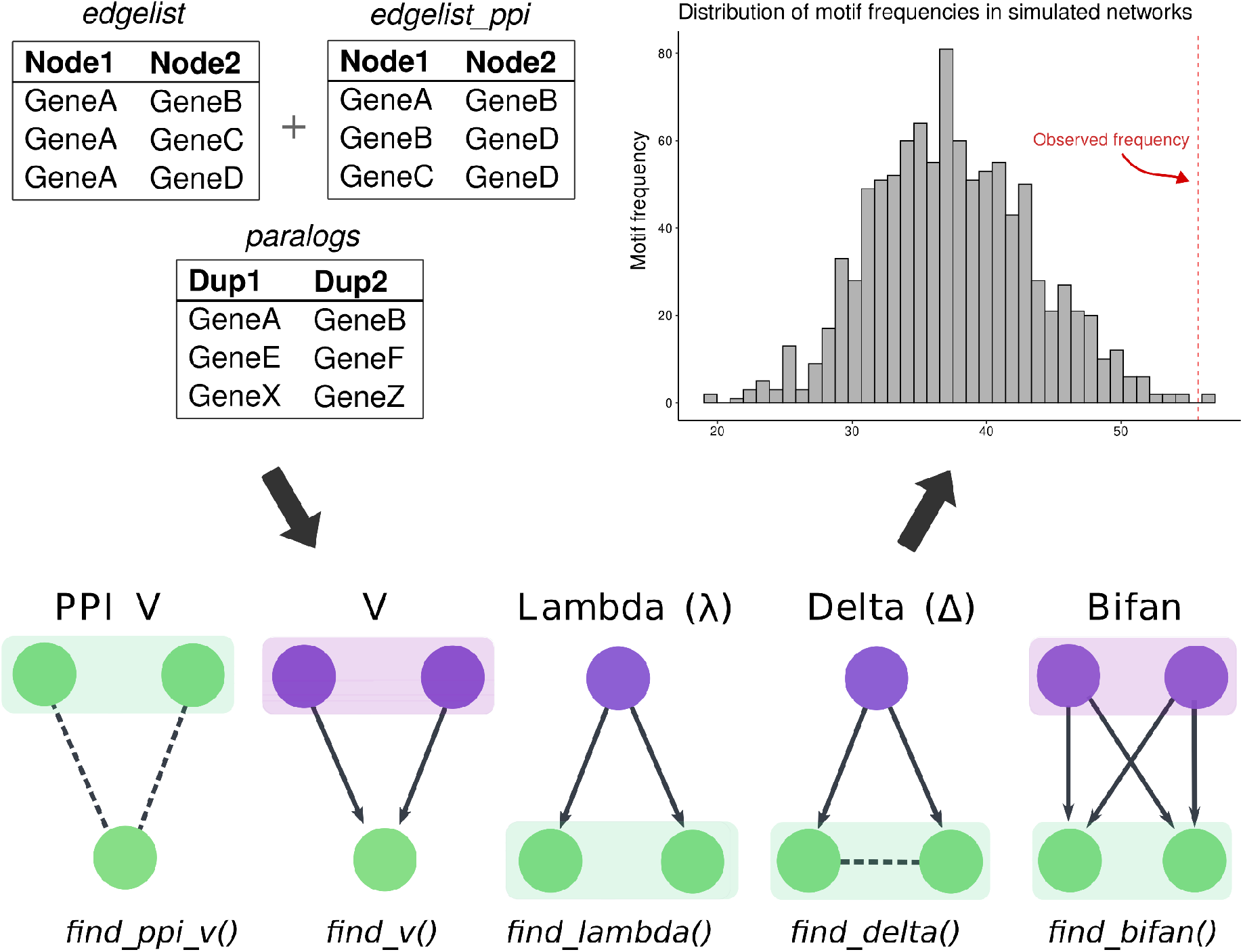
Workflow of motif identification and significance assessment with *magrene*. Input data types for motif detection, which include a data frame of GRN edges, a data frame of PPI network edges, and a data frame of paralogous gene pairs. Motif types and R functions in *magrene* to find specific motifs are indicated in the bottom panel. Shaded boxes indicate paralogous gene pairs. Regulators and targets are indicated in purple and green, respectively. Arrows indicate directed regulatory interactions, while dashed lines indicate protein-protein interactions. Observed motif frequencies for each motif type are then compared to a null distribution obtained from simulated networks, as indicated in the histogram. See Materials and Methods for details.

## Results and Discussion

### Paralogous gene pairs, summary statistics, and network topology

We identified and classified duplicated genes in the genomes of seven angiosperm species: *Glycine max, Arabidopsis thaliana, Solanum lycopersicum, Populus trichocarpa, Vitis vinifera, Zea mays*, and *Oryza sativa* (Table 1). These species were selected to account for a broad range of plant families and WGD histories. Expectedly, all species have more SSD-derived gene pairs than WGD-derived gene pairs, but such difference in frequency is smaller for species that have undergone recent WGD events (Table 1). Peaks in Ks distributions were used to ascribe duplicate gene pairs to their respective age groups (Table 1; Supplementary Figure S2, see Materials and Methods for details). All WGD and SSD comparisons presented hereafter were only performed with genes from the same age group, unless stated otherwise.

The degree distributions of all inferred GRNs and PPI networks satisfied the scale-free topology fit, indicating that they have the topological properties of real-world biological networks. As differences in motif frequencies can be due to differences in node degree, we compared the degree distributions of WGD- and SSD-derived genes in GRNs and PPI networks to detect potential biases. We observed no significant differences in degree distributions for WGD- and SSD-derived genes in both networks (Mann-Whitney U test; *P* < 0.05). Although some comparisons had significant differences (*e*.*g*., PPI network of *A. thaliana* and GRN of *G. max*), the small *P-*values are likely due to large sample sizes, as the effect sizes are negligible (rank-biserial correlation < 0.15) (Supplementary Figures S3 and S4).

### PPI networks are enriched in WGD-derived genes

We performed a Fisher’s exact test using all genes as background to test for associations between duplication modes and the presence of protein-protein interactions. We observed an enrichment of WGD-derived genes in the PPI networks for all species (*P* < 0.001), suggesting that strong selection pressures constrain the protein products of WGD-derived genes to interact physically. Functional enrichment analyses revealed that the set of interacting WGD-derived genes is enriched in genes associated with signal transduction, lipid metabolism, carbohydrate and amino acid oxidation, cell wall biogenesis, redox homeostasis, translation, and transcriptional regulation (Supplementary Table S2).

These findings are in line with the gene balance hypothesis, as interacting WGD-derived pairs are enriched in dosage sensitive biological processes, which tend to be retained more often after WGD events to preserve stoichiometry (Maere et al. 2005; Freeling and Thomas 2006; Freeling 2009; Birchler and Veitia 2010; Teufel et al. 2019; Almeida-Silva et al. 2020). Single-gene duplications for genes involved in complex PPI networks would likely be deleterious, as it leads to dosage imbalance. Thus, selection tends to retain such genes only if the whole PPI network is duplicated, as it happens in whole-genome duplication events (Maere et al. 2005).

### Selective pressures constrain sequence divergence in interacting WGD-derived genes

To understand how evolution shaped the functional divergence of WGD- and SSD-derived gene pairs, we explored how duplicates diverge in sequence over time. As synonymous substitutions accumulate in a neutral manner, Ks was used as a proxy for time, and Ka was used to represent sequence divergence. To account for saturation at higher Ks values, we fitted Michaelis-Menten curves to the scatter plot, as previously done by (Tasdighian et al. 2017; Defoort et al. 2019). For all species, we found that interacting WGD-derived gene pairs have slower rates of sequence divergence over time as compared to SSD-derived genes (Figure 2).

**Fig. 2.**
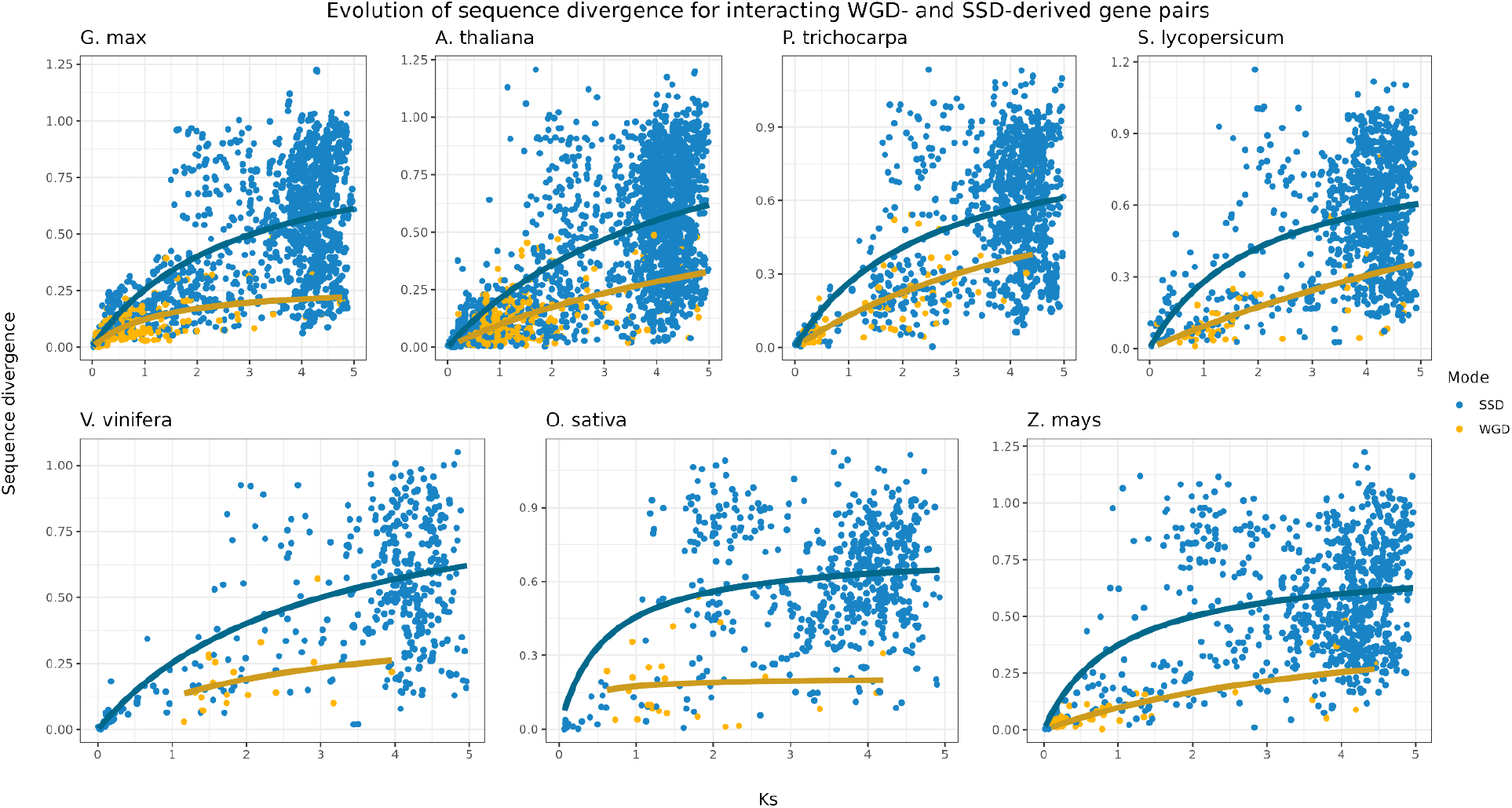
Evolution of sequence divergence for interacting WGD- and SSD-derived gene pairs. Rates of nonsynonymous substitutions per substitution site (Ka) were used to represent sequence divergence, and rates of synonymous substitutions per substitution site (Ks) were used as a proxy for time. As linear models cannot account for saturation at higher Ks values, a Michaelis-Menten curve was fitted in the scatter plot. The plot shows that WGD-derived gene pairs have lower sequence divergence over time as compared to SSD-derived pairs.

Sequence divergence leads to novel and/or specialized gene functions, which can disrupt gene balance. Thus, the lower sequence divergence for interacting WGD-derived genes is predicted by the gene balance hypothesis. Similar findings have been observed in *Arabidopsis thaliana* (Tasdighian et al. 2017; Defoort et al. 2019), *Saccharomyces cerevisiae* (Fares et al. 2013), and tomato and maize (Defoort et al. 2019). Collectively, these findings suggest that this is a universal pattern for WGD-derived genes, as it has been observed in multiple independent taxa and studies.

### WGD-derived gene pairs tend to interact with the same partners

For each species and age group, we calculated the interaction similarity of paralogous gene pairs using Sorensen-Dice indices, a metric that ranges from 0 to 1, where 0 means no shared partners, and 1 means complete overlap of partners. Overall, WGD-derived gene pairs have a higher interaction similarity in PPI networks than SSD-derived pairs (Figure 3). The difference is even more pronounced for older duplicate pairs, with ancient WGD-derived pairs displaying a much higher interaction similarity than ancient SSD-derived pairs (Figure 3). However, we observed no difference in interaction similarity between WGD- and SSD-derived pairs in GRNs (Supplementary Figures S5 and S6).

**Fig. 3.**
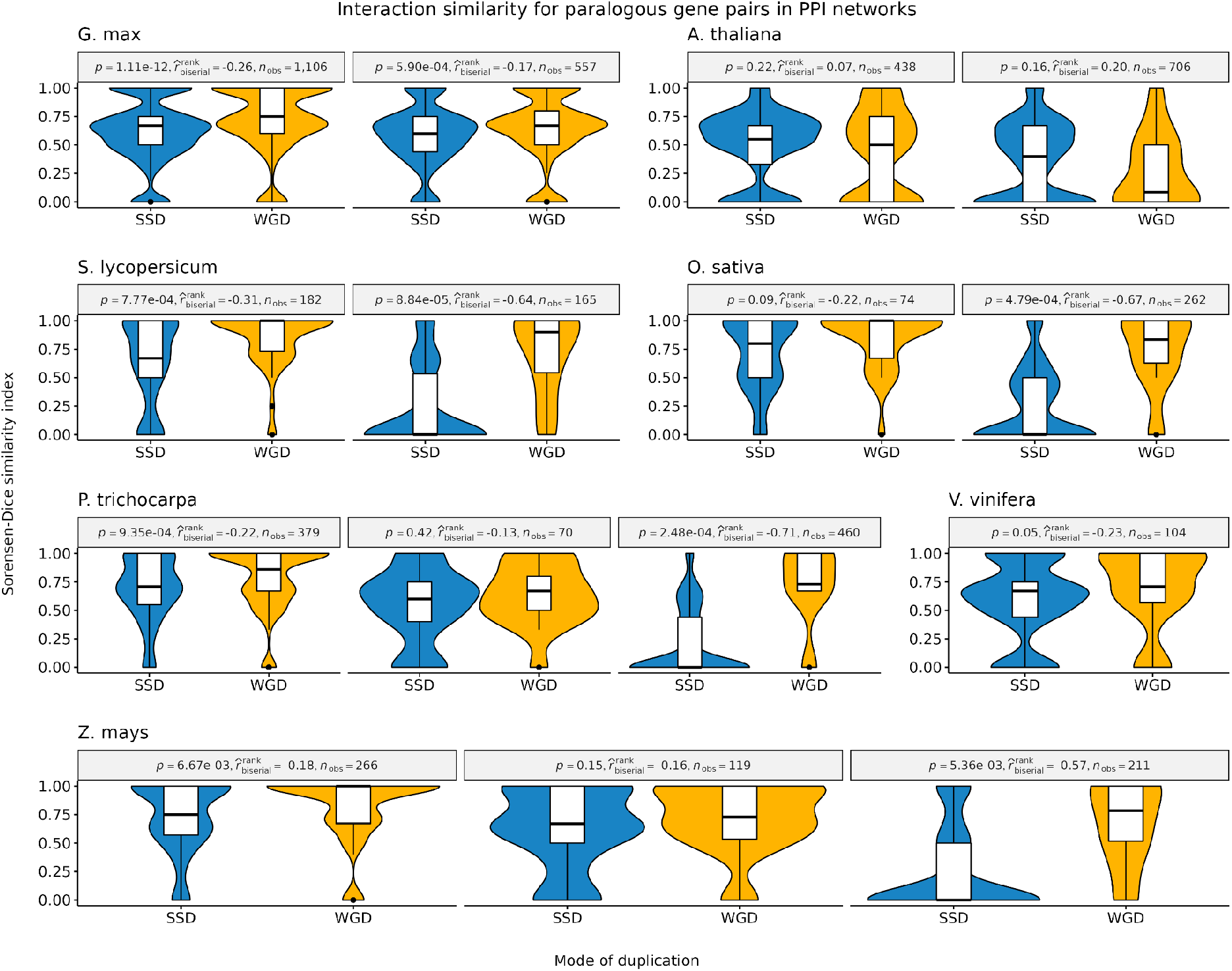
Interaction similarity for WGD- and SSD-derived gene pairs in PPI networks. Sorensen-Dice similarity indices were used to indicate interaction similarity. Overall, WGD-derived gene pairs have a higher interaction similarity as compared to SSD-derived gene pairs, especially for older duplicates (Mann-Whitney U test; P < 0.05). Effect sizes were measured using point-biserial correlation coefficients.

The observed patterns in PPI networks suggest that strong selection pressures constrain WGD-derived pairs to preserve their interactions over time, while more relaxed pressures allow SSD-derived pairs to lose and gain interaction partners. The higher preservation of interacting partners for WGD-derived gene pairs was previously observed in *A. thaliana* and human PPI networks (Defoort et al. 2019; Mottes et al. 2021), and it can again be explained by dosage balance constraints. As SSD-derived genes are generally not dosage sensitive, they can establish novel connections in PPI networks. On the other hand, novel connections for WGD-derived genes are likely disadvantageous, because they can disrupt the stoichiometric balance in the cell.

### (Recent) whole-genome duplications fuel(ed) the emergence of network motifs

We identified and counted network motifs for each species and age group using the R package *magrene* (see Methods). We also counted motif frequencies in 1,000 degree-preserving simulated networks, and Z-scores were used to assess the significance of observed motif frequencies. We found that genes derived from recent WGD events are more frequently part of network motifs than genes from more ancient WGD events, demonstrating that motifs are being lost over time (Figure 4A). We also found that species with recent WGD events generally have a higher motif frequency than species with more ancient WGDs, regardless of the duplication mode that created the genes forming motifs (e.g., *Z. mays* vs *O. sativa*; *G. max, A. thaliana, and P. trichocarpa vs V. vinifera* in Figure 4B).

**Fig. 4.**
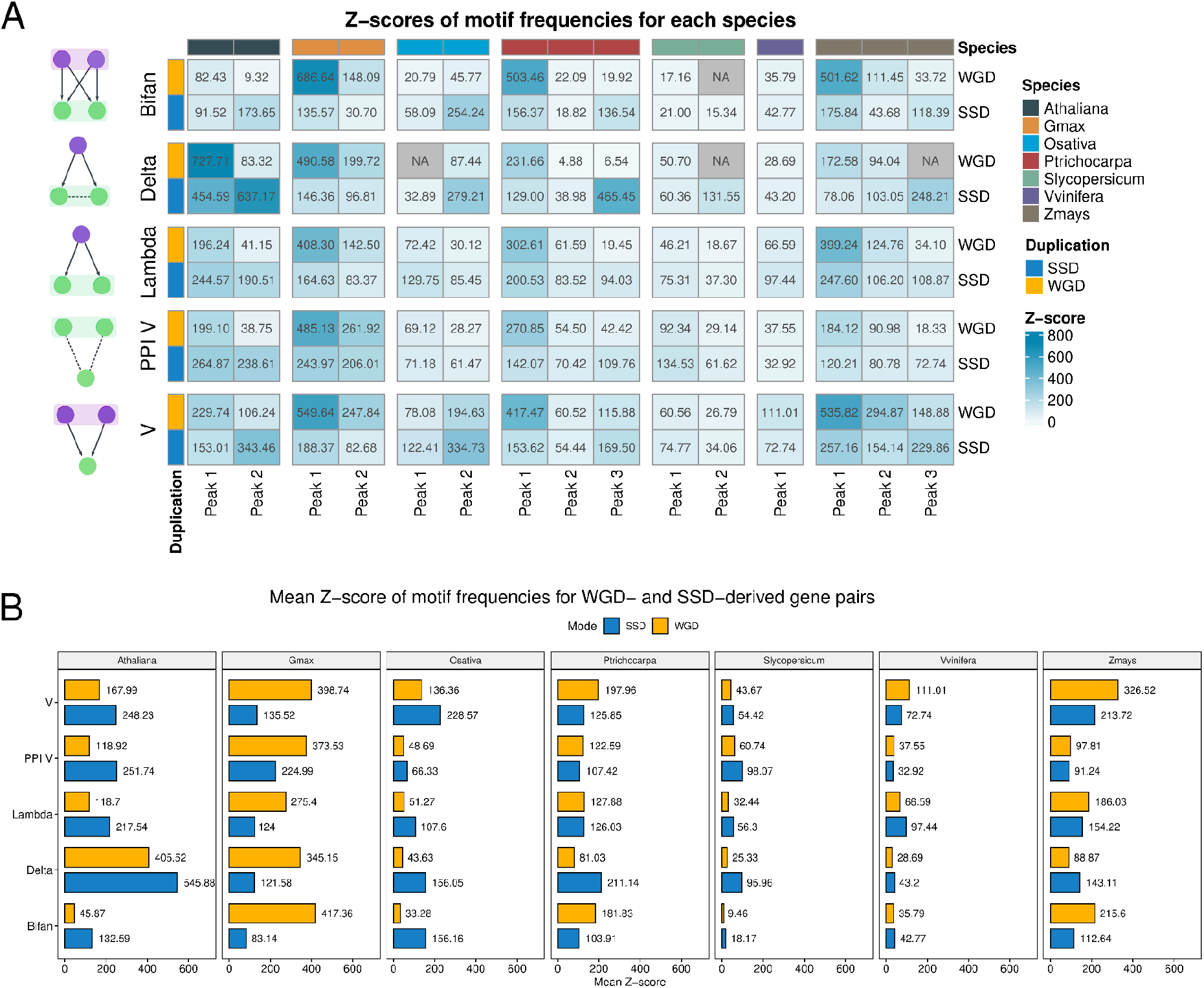
Z-scores of network motif frequencies in each species. A. Z-scores of motif frequencies by species and Ks peak-based age group. Z-scores <2 were considered not significant and set to NA. The plot shows that genes derived from recent WGD events are more frequent in network motifs than genes from more ancient WGD events. B. Mean Z-score of motif frequencies for WGD- and SSD-derived gene pairs. Recent polyploids have higher motif frequencies than ancient polyploids. WGD-derived genes have a greater contribution to motif formation in recent polyploids, while SSD-derived genes are more frequent in ancient polyploids.

While the *O. sativa* genome has signatures of a WGD event that is shared with all Poaceae, the *Z. mays* genome has undergone an additional WGD event ∼5-12 million years ago (Wei et al. 2007; Van de Peer et al. 2017). Likewise, the *V. vinifera* genome has signatures of a WGD event that is shared with all core eudicots, while the genomes of *P. trichocarpa, A. thaliana*, and *G. max* have undergone 1, 2, and 2 additional rounds of WGD, respectively (Van de Peer et al. 2017; Clark and Donoghue 2018). The higher motif frequencies in species with additional WGDs reveal that recent polyploidization events played a key role in the increase of robustness (*i*.*e*., greater number of nodes, edges, and interactions thereof) in gene regulatory networks, providing genomes with the raw genetic material (*i*.*e*., network nodes) for evolving novel regulatory interactions. However, it is noteworthy that such increased motif frequencies were not observed for *S. lycopersicum*, whose genome has signatures of an additional, Solanaceae-specific whole-genome triplication event (Figure 4A and 4B). This exceptional case suggests that rates of motif loss vary across clades, and *S. lycopersicum* (or even all Solanaceae) might have faster rates of motif loss.

In human GRNs, Mottes *et al*. (2021) observed that SSD-derived genes have a greater contribution to the formation of network motifs than WGD-derived genes. However, the two WGD events studied happened around the emergence of vertebrates (∼500 million years ago) (Van de Peer et al. 2009), hence they are very ancient events. Here, by analyzing WGD events of different ages, we demonstrate that the findings from Mottes *et al*. (2021) are only true for ancient, but not for recent WGD events. We observed that whether WGD or SSD has a greater contribution to motif formation is tightly linked to the WGD background of the species (*i*.*e*., ancient vs recent WGD). WGD-derived gene pairs have a greater contribution to motif formation in species with recent WGD events (e.g., *G. max* and *Z. mays*), while SSD-derived genes overtake WGD genes in motifs in species with more ancient WGD events (Figure 3B). However, it is not clear whether motifs are lost over time because genes involved in motifs are lost (as a result of fractionation), or because networks are rewired.

Our findings demonstrate that WGD and SSD events contributed in different ways to the evolution of gene regulatory networks in angiosperms, as recently observed for human gene regulatory networks (Mottes et al. 2021). The greater contribution of recent WGDs to motif formation suggests that WGD events have a more significant impact on the short-term evolution of polyploids. In line with our findings, polyploidy and the short-term redundancy it generates have been associated with an increased robustness against mutations and environment changes (Crow and Wagner 2005; Van de Peer et al. 2009). For instance, WGD events have been correlated with surviving environmental turmoil, such as during the Cretaceous-Paleogene extinction and glaciation events (Vanneste et al. 2014; Van de Peer et al. 2017; Novikova et al. 2018; Yao et al. 2019; Koenen et al. 2021). Assuming motif frequencies as a proxy for GRN robustness, the greatest chances of survival for recent polyploids could be explained by their increased GRN robustness. Alternatively, as motifs are genetic circuits that perform elementary regulatory functions, the greater number of motifs in recent polyploids creates bigger signals due to the combinatorial effect of each individual motif. Ultimately, such increased signals would allow greater jumps in the fitness landscape that would not be possible with small-scale duplications, possibly explaining an increased chance of survival during periods of extinction.

### Network motifs created by WGD and SSD are associated with different biological processes

To explore how WGD and SSD shaped angiosperm GRNs from a functional perspective, we performed functional enrichment analyses for GO terms, Interpro domains, and TF families for genes in motifs. Only bifans were enriched in particular functional categories or TF families. Bifans are combinatorial decision-making circuits that integrate different input signals, organizing the transcription of downstream target genes. We found that WGD and SSD contribute to GRNs with genes of different functional groups. WGD-derived genes in bifans are enriched in genes associated with dosage-sensitive processes, while SSD-derived genes in bifans are enriched in stress-related processes (Figure 5; Supplementary Table).

**Fig. 5.**
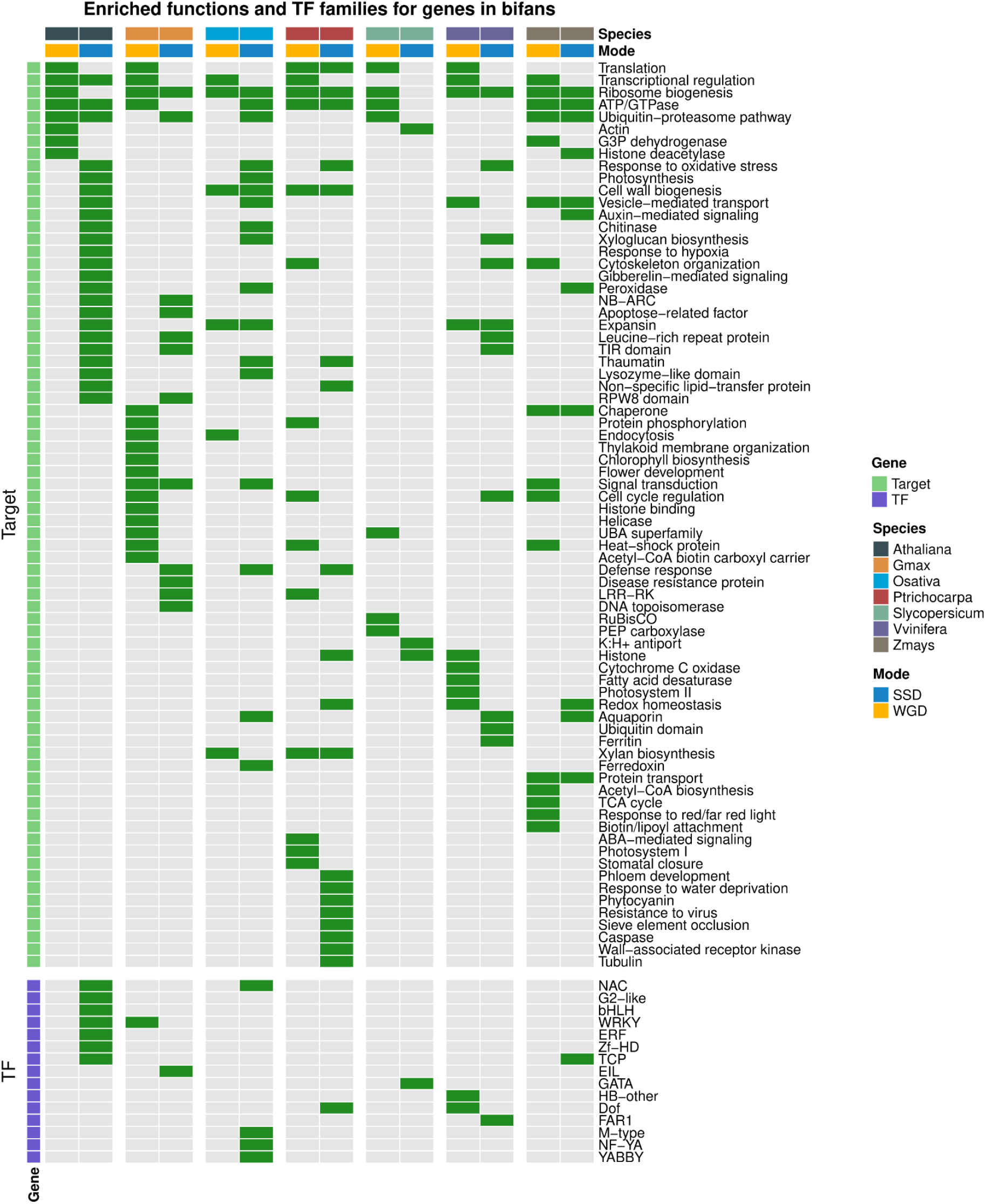
Enrichment of transcription factor families and gene functions for WGD- and SSD-derived genes in bifans. Enrichment analyses for TF families in V and bifans were performed using all TFs in the GRN as background. Green and gray cells represent enrichment and no enrichment, respectively. Enrichment analyses for functional categories in lambda, delta, and bifans were performed using all targets in the GRN as background. WGD-derived genes in bifans are enriched in dosage-sensitive processes, such as transcriptional regulation, regulation of cell cycle, translation, and photosynthesis, while SSD-derived genes in bifans are enriched in stress-related processes, such as response to hypoxia and oxidative stress, pathogenesis-related proteins, recognition of pathogen-associated molecular patterns, and stress-related TF families, such as ERF, NAC, and WRKY.

WGD-derived bifans are enriched in genes associated with translation, transcriptional regulation, flower development, histone modifications, photosynthesis-related processes (*e*.*g*., chlorophyll biosynthesis, thylakoid membrane organization, and RuBisCO), cell cycle regulation, and carbohydrate and lipid metabolism (Figure 5; Supplementary Table S3). The association of WGD-derived genes with such functional categories has been reported in multiple independent studies (Sato et al. 2009; Conant et al. 2014; Freeling et al. 2015; Almeida-Silva et al. 2020), and it is likely because these gene families are more dosage sensitive than others (Cheng et al. 2018).

The enriched stress-related processes for SSD-derived target genes in bifans include oxidative stress, response to hypoxia, pathogenesis-related proteins (*e*.*g*., chitinases and lysozymes, peroxidases, and thaumatins), recognition of pathogen-associated molecular patterns (e.g., leucine-rich repeat receptor kinase and wall-associated kinases). Likewise, SSD-derived TFs in bifans are enriched in stress-related TF families, such as WRKY, ERF, NAC, and G2-like. Previous studies have reported that stress-related gene families tend to expand through tandem duplications, because their organization in tandem arrays facilitates their coexpression in response to environmental stimuli (Amoutzias and Van de Peer 2008; Hanada et al. 2008; Rodgers-Melnick et al. 2012). Although WGD is the main force driving the expansion of TFs due to dosage balance, hence increasing the number of nodes in GRNs, our findings demonstrate that SSD also plays an important role in GRNs by contributing with stress-related genes.

Collectively, the patterns we observe for WGD- and SSD-derived motifs in this study are very similar to what has been observed for WGD- and SSD-derived individual genes. In summary, we show that: i) WGD-derived genes and motifs are lost over time, but some are retained over long evolutionary timescales; ii) WGD-derived genes and motifs that are retained long-term are associated with growth and development, with a particular preferential retention of dosage-sensitive molecular pathways, such as transcriptional regulation, histone modifications, and cell cycle regulation; and iii) SSD-derived genes and motifs that are retained long(er)-term are associated with stress-related functions. Such similarities suggest that the evolution of motifs is intricately associated with the evolution of genes, and a more detailed picture could be observed by comparing rates of gene and motif loss.

For instance, we observed that WGD-derived motifs are quickly lost over time as compared with SSD-derived motifs (see previous section, Fig. 4a). On the one hand, the rapid loss of WGD-derived motifs could be explained by the rapid loss of WGD-derived genes due to fractionation (Freeling et al. 2015). On the other hand, if rates of motif loss were higher than rates of gene loss, it would suggest that motifs are lost first (*i*.*e*., due to transcriptional rewiring), leading to relaxed selection pressures that could eventually lead to gene loss or, in special cases, to subfunctionalization and/or neofunctionalization. The latter hypothesis implies that participating in motifs imposes additional selective constraints on genes and, hence, motifs could have a significant long-term impact on evolution.

Additionally, as motifs are subgraphs that are observed more often than is expected by chance, they are thought to have been positively selected (Mottes et al. 2021). We hypothesize that selection acting upon motifs leads to a higher chance of retention for genes that participate thereof. In the long run, transcriptional rewiring of motifs created by WGD events could contribute to evolutionary innovations and increase in organismal complexity, such as what has been observed for the auxin response regulators AUX/IAA and some subfamilies of MADS-box transcription factors in plants (Remington et al. 2004; Veron et al. 2007), and the RAR/RXR pathway and homeobox (Hox) gene clusters in vertebrates (Holland et al. 2008; Mottes et al. 2021).

## Conclusions

In this paper, we explored the impact of WGD and SSD events on the topology of gene regulatory networks in angiosperms. We observed that each duplication mechanism contributed to network evolution in its own way. Using a diverse set of species with different numbers and ages of WGD, we confirm previous observations that WGD-derived genes are under strong selective constraints, have slower rates of sequence divergence, and tend to preserve interactions with themselves and with other partners. These findings provide robust evidence that previously reported patterns for WGD-derived genes are universal, regardless of the number and age of genome duplications. WGD events contribute to GRNs with genes associated with dosage-sensitive processes, while SSD contributes with stress-related genes. Strikingly, recent polyploids have higher motif frequencies than ancient polyploids, suggesting that polyploidy might have a great(er) impact on the short term. We hypothesize that WGD events confer a short-term selective advantage to polyploids by enhancing GRN robustness and potentially magnifying GRN output signals (e.g., traits, gene expression, etc.) through the combinatorial effect of each individual motif, resulting in greater jumps in the fitness landscape and increased chances of survival during periods of environmental instability, including mass extinction events. Our findings underscore the importance of WGD events in the evolution of GRNs and highlight their potential role in facilitating adaptation to changing environments.

## Acknowledgements

YVdP acknowledges funding from the European Research Council (ERC) under the European Union’s Horizon 2020 research and innovation program (No. 833522). YVdP and FA-S acknowledge funding from Ghent University (Methusalem funding, BOF.MET.2021.0005.01).

## Data availability

All data and code used in this paper are available in a GitHub repository (https://github.com/almeidasilvaf/polyploid_GRNs) to ensure full reproducibility.

## Notes

### Competing Interest Statement

The authors have declared no competing interest.

