## Supplementary Figures for "Whole-genome duplications and the long-term evolution of gene regulatory networks in angiosperms"

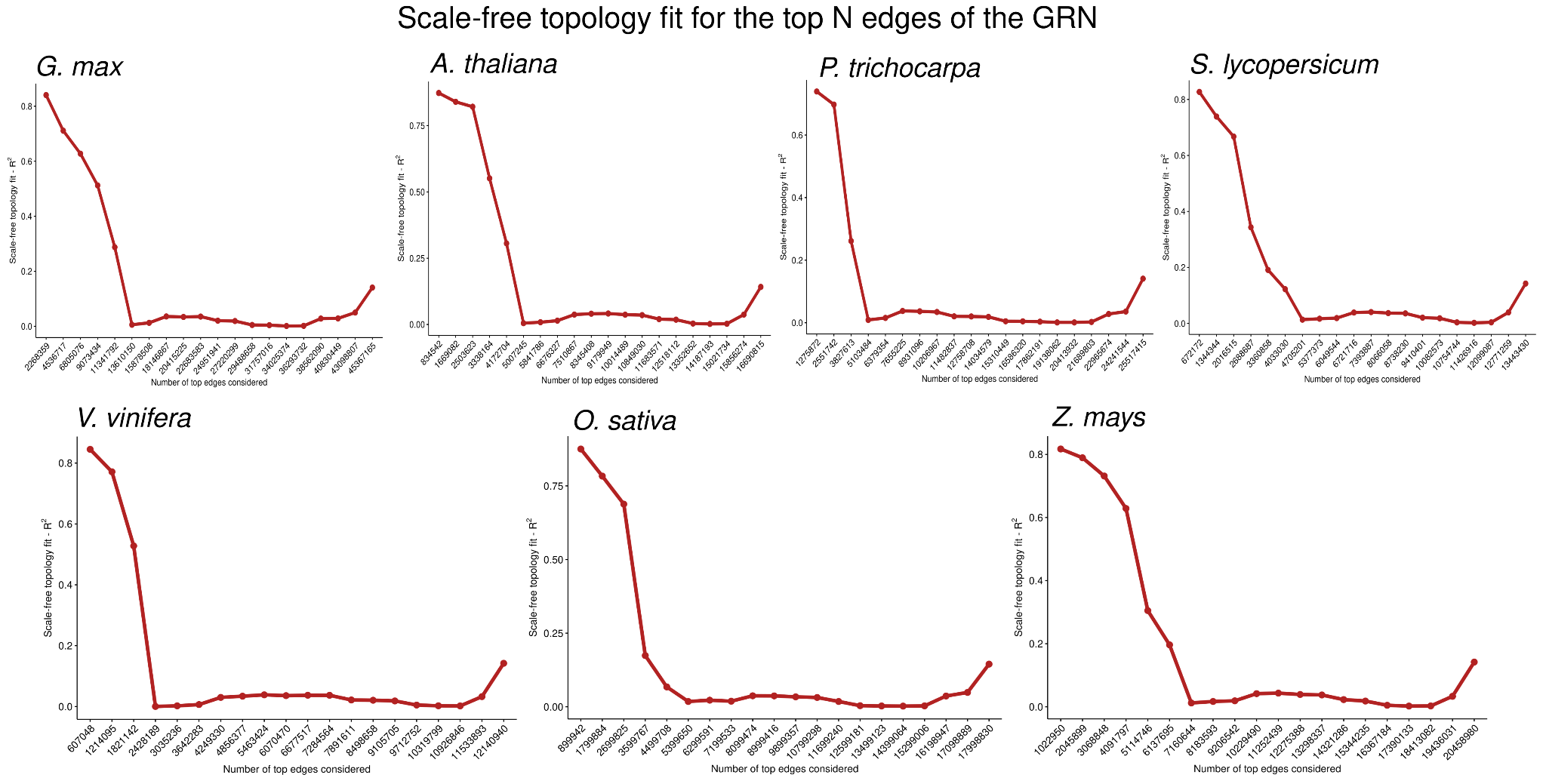

**Fig. S1. Scale-free topology fit for each subset of the fully connected gene regulatory networks**. For each species, subnetworks were created by ranking and extracting the top *N* edges (x axis). The original fully connected graph for each species does not satisfy the scale-free topology fit, but subsets of the top-ranked edges do. The highest value of *N* for which the network satisfied the scale-free topology fit (R^2^ = 0.75) was selected as optimal.

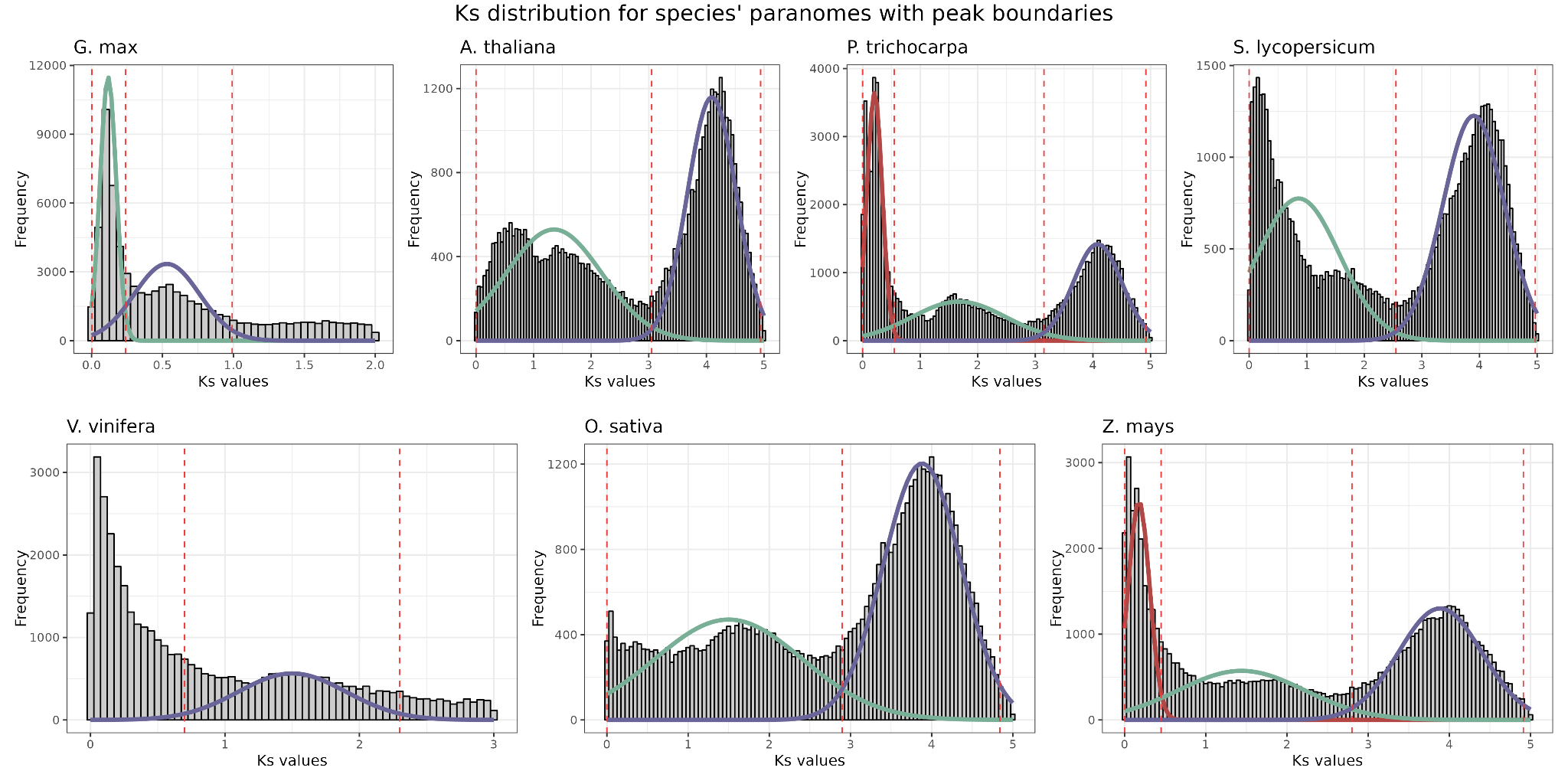

**Fig. S2. Peaks in Ks distributions for each species and age boundaries.** Density lines represent Ks peaks identified with Gaussian mixture models. Dashed red lines represent peak-based age boundaries, which were used to split duplicated gene pairs in age groups. Only WGD- and SSD-derived gene pairs from the same age group were compared, as age can be a confounder when comparing motif frequencies.

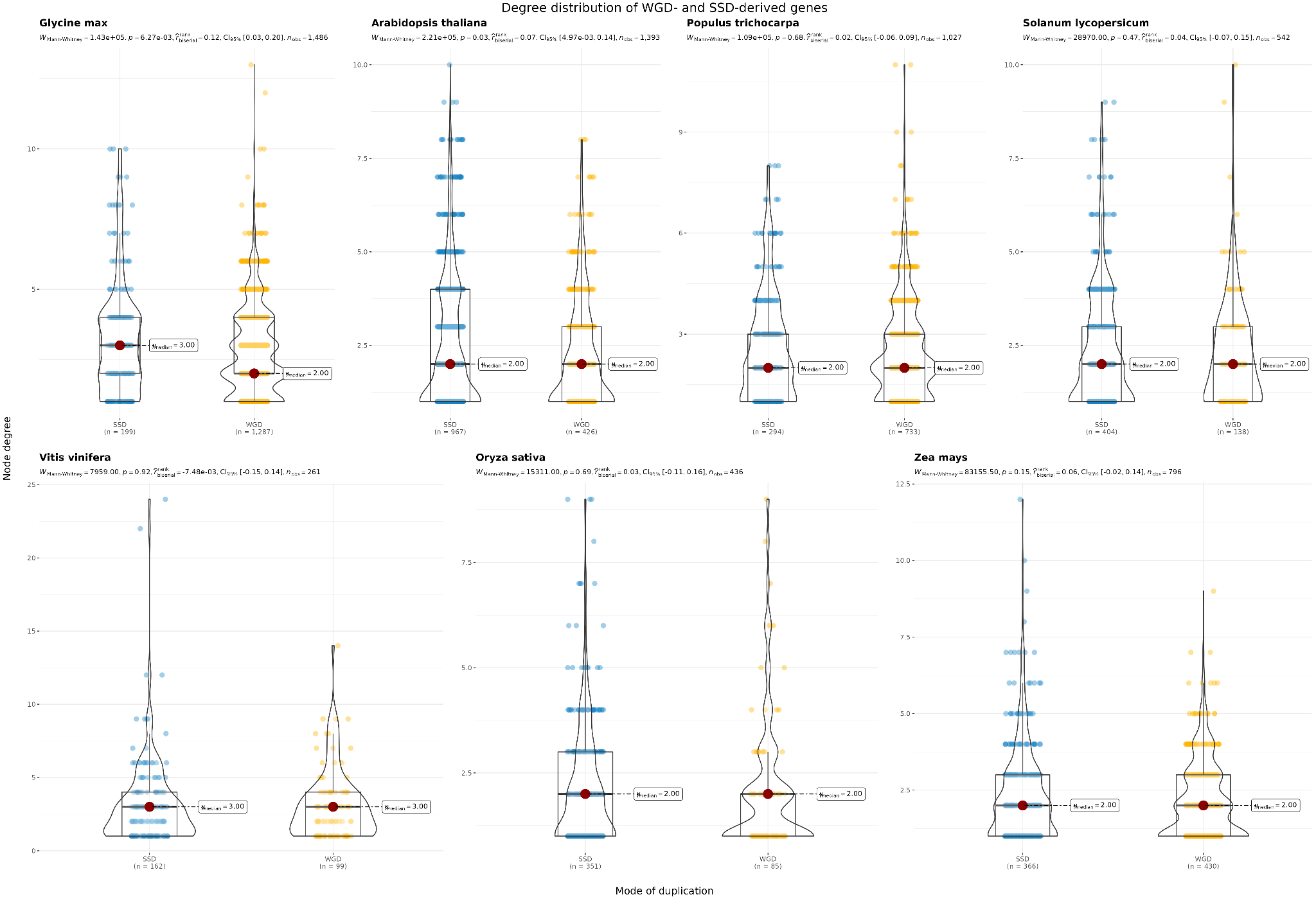

**Fig. S3. Comparison of the degree distributions for WGD- and SSD-derived genes in PPI networks.** The Mann-Whitney U test revealed no differences in degree distributions. Although some comparisons showed significant differences (*P* < 0.05), the effect size is negligible (rank-biserial correlation < 0.15), suggesting that the low *P-*values are likely an artifact resulting from large sample sizes.

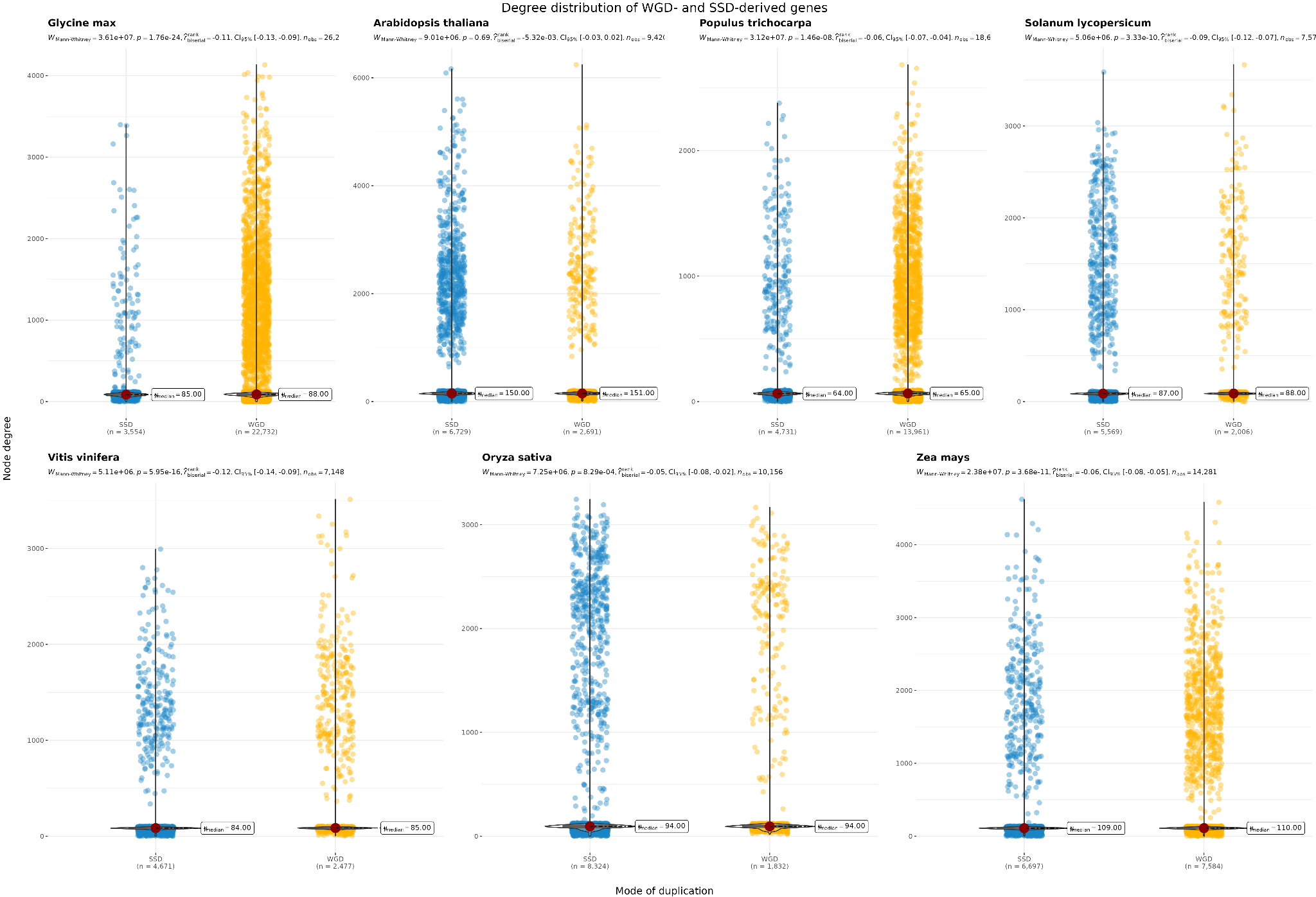

**Fig. S4. Comparison of the degree distributions for WGD- and SSD-derived genes in GRNs.** There were no differences in degree distributions (Mann-Whitney U test). Although some comparisons showed significant differences (*P* < 0.05), the effect size is negligible (rank-biserial correlation < 0.15), suggesting that the low *P-*values are likely an artifact resulting from large sample sizes.

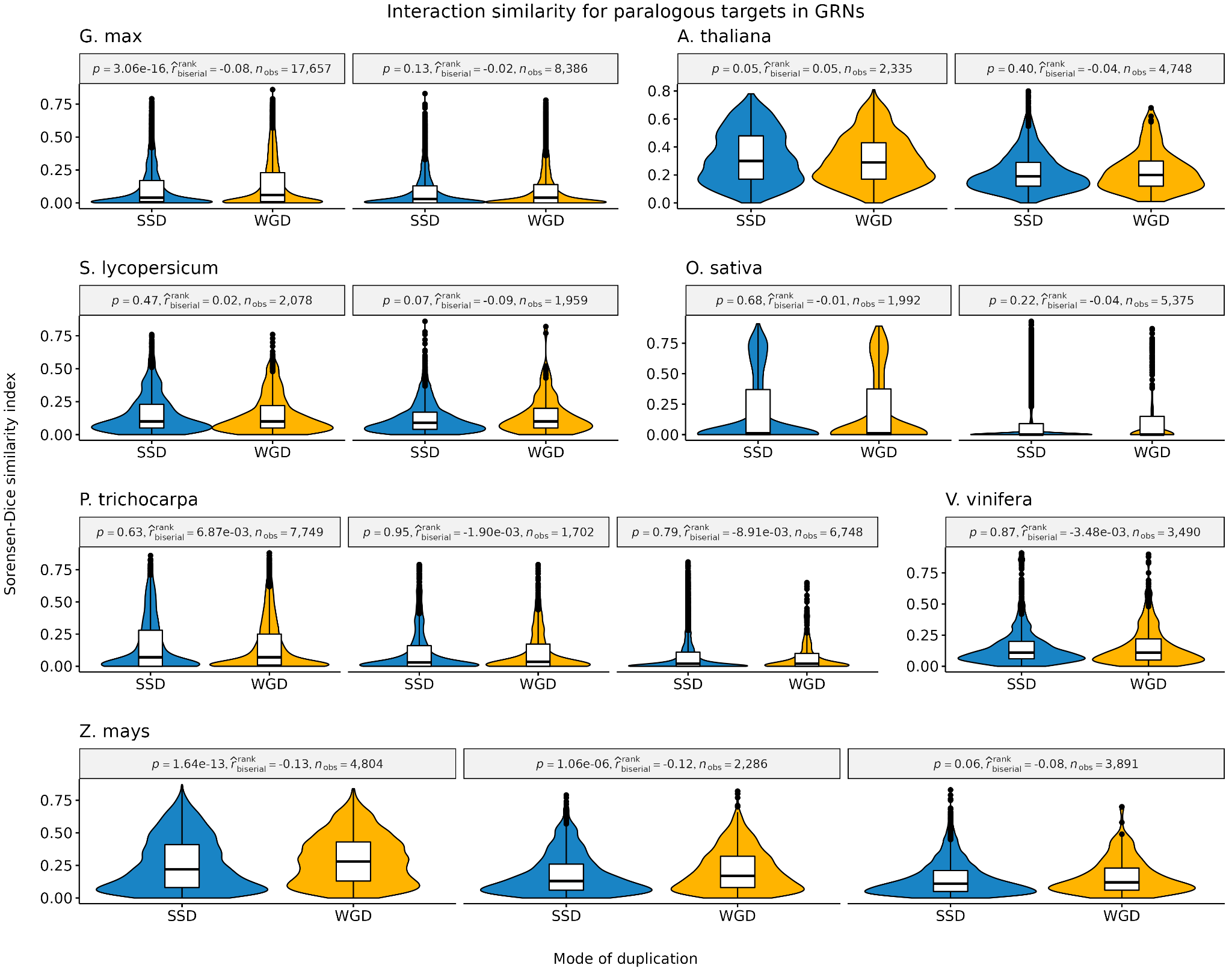

**Fig. S5. Interaction similarity between paralogous target genes in GRNs.** Sorensen-Dice similarity indices were used to indicate interaction similarity. No differences were observed between WGD- and SSD-derived gene pairs (Mann-Whitney U test; P < 0.05). Although some comparisons had significant *P*-values, it is likely an artifact resulting from large sample sizes, as effect sizes are negligible.

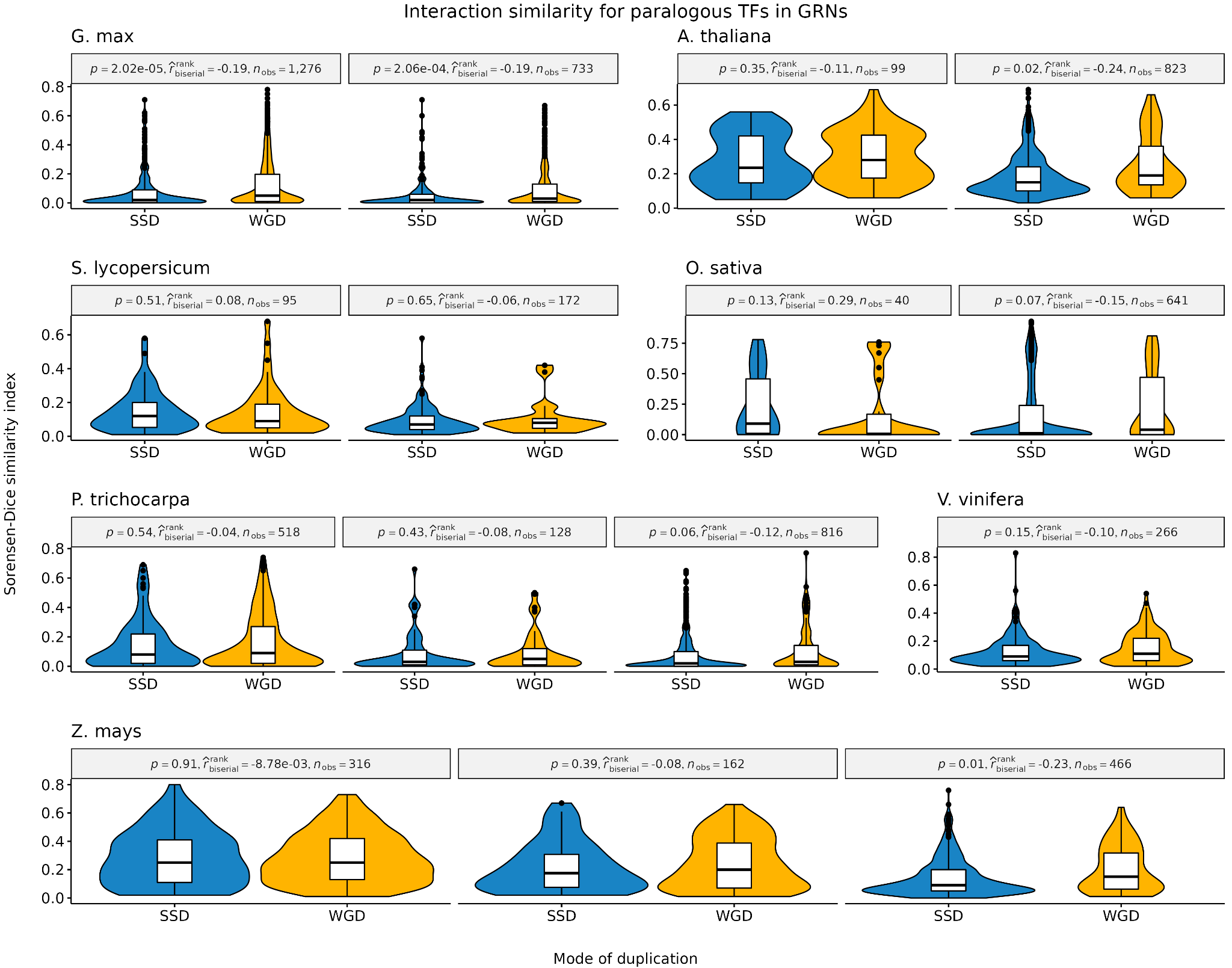

**Fig. S6. Interaction similarity between paralogous TFs in GRNs.** Sorensen-Dice similarity indices were used to indicate interaction similarity. Overall, no differences were observed between WGD- and SSD-derived gene pairs (Mann-Whitney U test; P < 0.05). Although some comparisons had significant *P*-values, it is likely an artifact resulting from large sample sizes, as effect sizes are negligible.
